# Benchmarking nearest neighbor retrieval of zebra finch vocalizations across development

**DOI:** 10.1101/2023.09.04.555475

**Authors:** Tomas Tomka, Xinyu Hao, Aoxue Miao, Kanghwi Lee, Maris Basha, Stefan Reimann, Anja T. Zai, Richard H. R. Hahnloser

## Abstract

Vocalizations are highly specialized motor gestures that regulate social interactions. The reliable detection of vocalizations from raw streams of microphone data remains an open problem even in research on widely studied animals such as the zebra finch. A promising method for finding vocal samples from potentially few labelled examples (templates) is nearest neighbor retrieval, but this method has never been extensively tested on vocal segmentation tasks. We retrieve zebra finch vocalizations as neighbors of each other in the sound spectrogram space. Based on merely 50 templates, we find excellent retrieval performance in adults (F1 score of 0.93 *±* 0.07) but not in juveniles (F1 score of 0.64 *±* 0.18), presumably due to the larger vocal variability of the latter. The performance in juveniles improves when retrieval is based on fixed-size template slices (F1 score of 0.72 *±* 0.10) instead of entire templates. Among the several distance metrics we tested such as the cosine and the Euclidean distance, we find that the Spearman distance largely outperforms all others. We release our expert-curated dataset of more than 50’000 zebra finch vocal segments, which will enable training of data-hungry machine-learning approaches.

## I. INTRODUCTION

In many species including humans, vocalizations play important roles during social behaviors such as aggressions, mating, breeding, and feeding. Inferring the functions of the vocalizations is a challenging task where machine learning could be promising^1^. The longitudinal study of vocalizations involves the challenging task of segmenting vocalizations from background noise. In vocal learners such as the zebra finch, the vocal segmentation task is particularly difficult, because the zebra finch vocal repertoire dramatically changes over the course of development^2,3^. Songs in young zebra finches start out as unstructured subsongs that lack categorical structure and that gradually differentiate into distinct classes of stereotyped syllables^4^. Zebra finches also produce less stereotyped calls^5^ with acoustic features that vary depending on behavioral context^5,6^.

To segment vocalizations in large vocal data sets, there is a growing literature on machine-learning based systems^7–10^. However, these systems have only recently been emerging and their potential is far from being fully explored. Foremost, for segmentation systems to perform well, they must be trained and tested on datasets of precisely segmented vocalizations. But to our knowledge, only one such dataset is publicly available^7,11^ and it contains merely 473 song syllables produced by a single adult male zebra finch and fails to include all vocalization types, so represents a biased sample of vocal output. Entirely lacking are public datasets of precisely segmented subsongs; a recent massive-data study on this important developmental phase^12^ simply ignores the segmentation problem and takes as proxy of vocalizations all amplitude-thresholded sound segments, semi-automatically excluding false positives in such a way to introduce false negatives (see Appendix). Unfortunately, amplitude thresholding can create severe problems if the recording quality is low^13^, which only emphasizes that this severe lack of training and test data forms a bottleneck for progress in large-scale research on vocal development, and it calls for the creation of gold-standard data sets.

One method for bootstrapping large vocal data sets from few precisely labelled samples is nearest neighbor (NN) retrieval^13^. NN retrieval is a highly successful information retrieval method^14^: it is used in tasks such as tagging images^15^, web mining^16^, recommendation systems^17,18^, and for inference in language models^19,20^. Although the computational cost of NN retrieval grows linearly with the number of templates and the size of the test recordings, NN search scalability has improved massively since the popularization of graphics processing units (GPUs) for parallel computing^21^ and with the advent of powerful approximate nearest neighbor methods^22–25^. One of the advantages of NN retrieval over neural networks is that NN retrieval uses few parameters and is interpretable^26–28^.

NN retrieval has been applied previously to the problem of birdsong analysis^29,30^. Brooker and colleagues used Pearson-correlation-based NN retrieval to benchmark commercially available song detection software such as MonitoR^30,31^. Anderson and colleagues even applied a dynamic time-warping algorithm to find data frames in the search space based on their minimal path-traversing distance to template frames^29^. However, the sample sizes and scopes of these works are very restrictive: they are based on single birds and unique distance measures^29^ and they excluded certain vocalization types from the analysis^30^.

We set out to scale up NN retrieval methods for annotating and proofreading vocal segments. The segmentation task we consider is to determine for each time point in a sound spectrogram (i.e., 16-ms sound interval) whether it contains a vocalization or not. We benchmark the performance of our approaches on two data subsets of adult (Subset 1) and juvenile (Subset 2) male zebra finch vocalizations. In our WHOLE approach, we use entire templates for NN retrieval, whereas, in the PART approach, we use fixed windows cut from the templates. The PART approach allows the detection of vocalizations from conserved parts and offers the practical benefit of yielding samples of fixed dimensionality. Among the many spectrogram-based distance metrics we apply during retrieval, we find that the Spearman distance outperforms all other metrics. We release our gold standard (GS) data set of more than 50’000 annotations, taking care of eliminating false negatives, i.e. vocalizations buried in noise that are easily missed by inattentive annotators.

## II. METHODS

### A. Sound recordings and spectrograms

We used data sets from four adult and four juvenile male zebra finches (each of the latter was recorded at three different ages, see Table I for details). Recording was triggered by vocalizations (or other sounds); thus, recordings are unevenly spaced in time depending on the activity of the bird. Each recording/file contains vocalizations with some silence before and after the vocalizations.

All adult birds (Subset 1) were raised in the animal facility of the University of Zurich. During recording, birds were housed in single cages in custom made soundproof recording chambers equipped with a wall microphone (Audio-Technica Pro42), and a loudspeaker. The day/night cycle was 14*/*10 h. Vocalizations were saved using custom song-recording software (Labview, National Instruments Inc.). Sounds were recorded with a wall-attached microphone and were digitized at 32 kHz. We analyzed data from birds that had already spent at least three days in their cage.

Data from juvenile birds (Subset 2) were randomly sampled from a publication^32^: We randomly selected 4 birds and from each bird we selected 3 days. Sounds in^32^ were recorded at a sampling rate of 44.1 kHz.

We computed sound spectrograms by Fourier transforming sound segments *X_t_ ∈* ℝ*^b^* of b= 512 samples. Accordingly, a spectrogram column *Y_t_ ∈* ℕ*^b^* at time *t* is given by Eq. (1), where Ω is a hamming window of length b= 512, and *β* = 6.54 for Subset 1 and *β* = 4.93 for Subset 2 is a parameter that controls the dynamic range of the int8 down conversion.

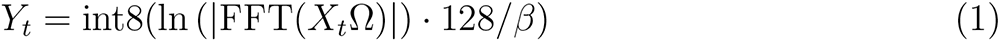

The hop size Δ*t* between adjacent Fourier segments is 128 samples corresponding to 4 ms in adults. For distance computations, we removed low frequencies (0-688 Hz in adults and 0-947 Hz in juveniles) due to the large background noise in these ranges.

### B. Generation of gold-standard annotations

From each day-long recording, we annotated a subset of data by randomly selecting a set of files. We annotated vocal segments (not further classified into vocalization types) with high temporal accuracy. To generate these gold-standard (GS) annotations, we used a semi-supervised segmentation method^13^, correcting poor segments and eliminating false positives by visual inspection of spectrograms. To eliminate false negatives, the present NN method was used with the cosine distance as metric. The GS dataset contains a label for each spectrogram column (“1” for vocal, and “0” for non-vocal). A detailed annotation protocol is provided in the “Supplementary information”.

### C. Nearest neighbor vocalization retrieval using gold-standard templates

A simple approach to retrieving sounds segments corresponding to vocalizations is to take a single template vocalization of (whole) duration *τ* and to compute spectrogram-based distances to all candidate segments from the search space. Candidates are contained in spectrogram windows of the same duration *τ* . The best candidate segment is the one with minimal spectrogram-distance to the template and that does not temporally overlap with the template, Fig. 1. To reduce computational cost, we restricted the search space to non-silent periods (defined by thresholding the root-mean-squared audio signal) of duration *≥ τ* .

When many templates are given, we generalize this single-template procedure to many templates by iteratively retrieving the top segments one-by-one, as described in the following.

**FIG. 1.**
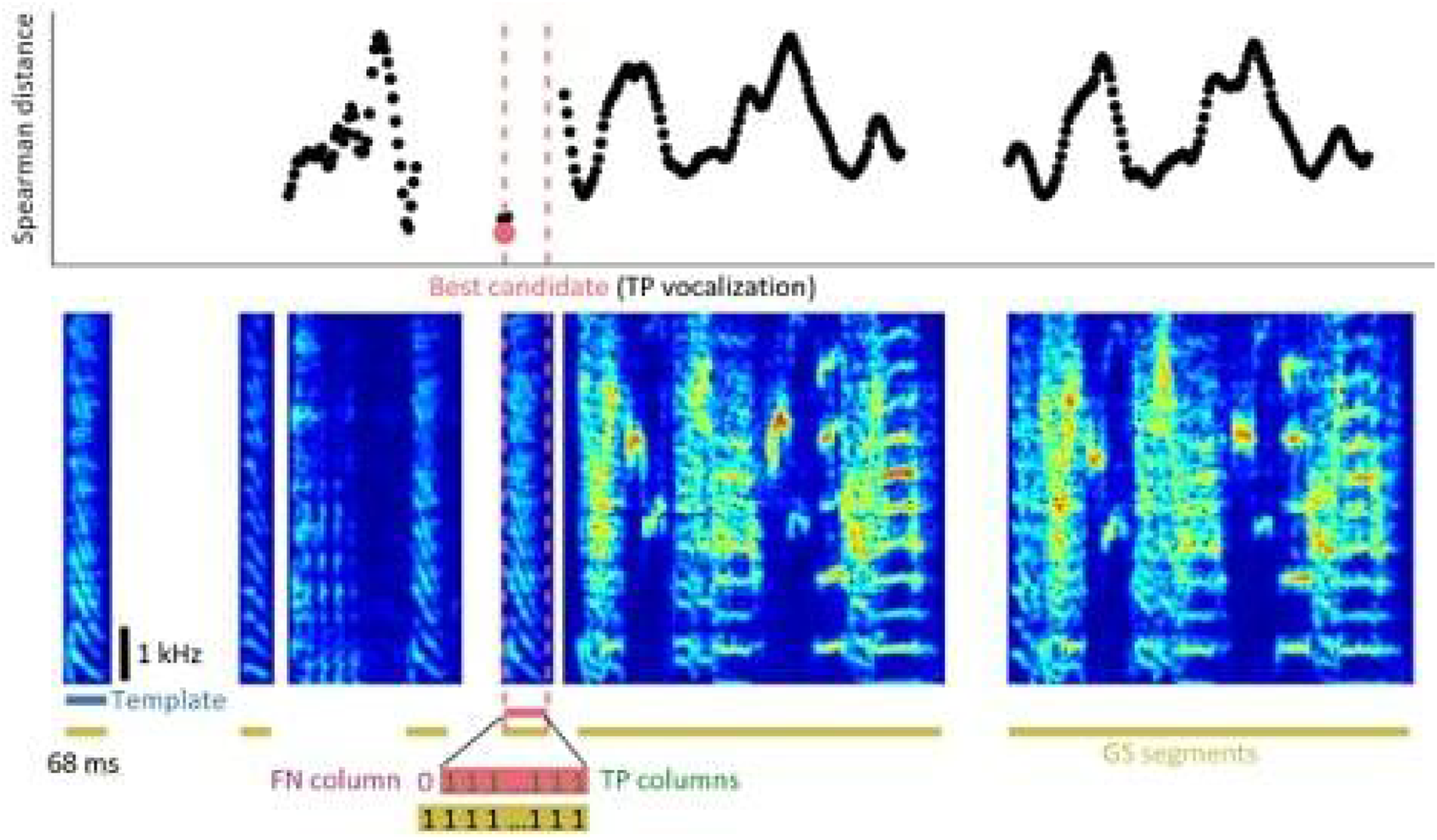
Template-based nearest-neighbor (NN) retrieval of vocal segments (WHOLE approach). For an exemplary template (leftmost spectrogram) drawn from our gold-standard (GS) dataset, we plot the (here Spearman) distance (top, dots aligned to candidate onsets) to all candidate segments of the same duration within the search space (other spectrograms). The best candidate (delimited by red dashed lines) is the one with minimal spectrogram-based distance (red dot, top). With this procedure, segmentation errors can arise from mismatching segment durations. Here, the best candidate starts one spectrogram column too late relative to the GS segmentation, giving rise to a false negative (FN) spectrogram column (purple 0). Since this error is within a reasonable tolerance (*≤* 5 columns), we regard this vocal segment (red horizontal bar) as containing a true positive (TP) vocalization.

### D. Vocalization retrieval using WHOLE approach

In the WHOLE approach (Fig. 2),we computed the spectrogram-based distances *D_ij_* of all template-candidate pairs. The distance *D_ij_* represents the distance between the *i − th* template (*i* = 1*, . . ., M*) and the *j − th* candidate in the search space. For a given template i, the search space is given by the set of candidates of the same duration *τ_i_* as the template. After we computed all distance pairs, we identified the best candidate segment to any template as the one with minimal distance, *argmin D_ij_*. After choosing the best segment, we removed it from the search space, thereby also removing candidates that overlapped with the best segment. Then we selected the next-best segment in an iterative procedure. By iteratively selecting the segment with minimal distance to any template, we chose a very greedy strategy of retrieving segments from the set of templates. In practice, we first computed all pairwise distances and maintained an index of valid candidate-template pairs to avoid re-computing any distances during the iterative procedure.

Because templates are of different durations *τ_i_*, they might bias this retrieval process to short templates. To address this possibility, we tested four different normalizations of distances: no normalization, dividing distances *D_ij_* by *τ_i_*, by 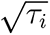, or min-max normalizing them for each template separately as in Eq.(2).

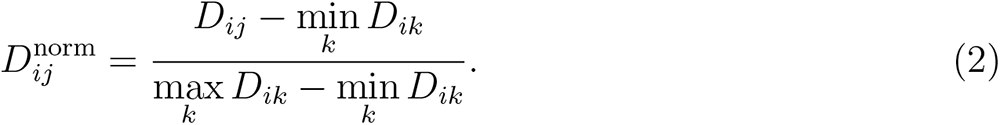

### E. Vocalization retrieval using PART approach

In the PART approach, we circumvent any duration-induced distance bias by slicing each template into overlapping slices of w spectrogram columns (Fig. 2), where the integer parameter w is shorter than a typical template. To any template i with duration *τ_i_ < w*, we appended a trailing zero-pad so that all templates had a duration of at least w. From M templates, we obtained in total 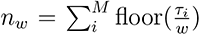 template slices. We then computed all distance pairs *D_ij_* between template slices and candidate slices. We then chose the best candidate slice as the one with minimal distance to any of the *n_w_* template slices. Based on the best candidate slice, we selected the associated best segment as the sound interval with the same relative timing as the template the slice was taken from (the onset and offset of the best segment formed the same time lags to the slice as did the onset and offset of the sliced template), Fig. 2. Thus, the best candidate segment was selected to be of equal duration as the sliced template. There was one exception to this procedure: when the selected best segment extended into a silent period, it was cropped.

**FIG. 2.**
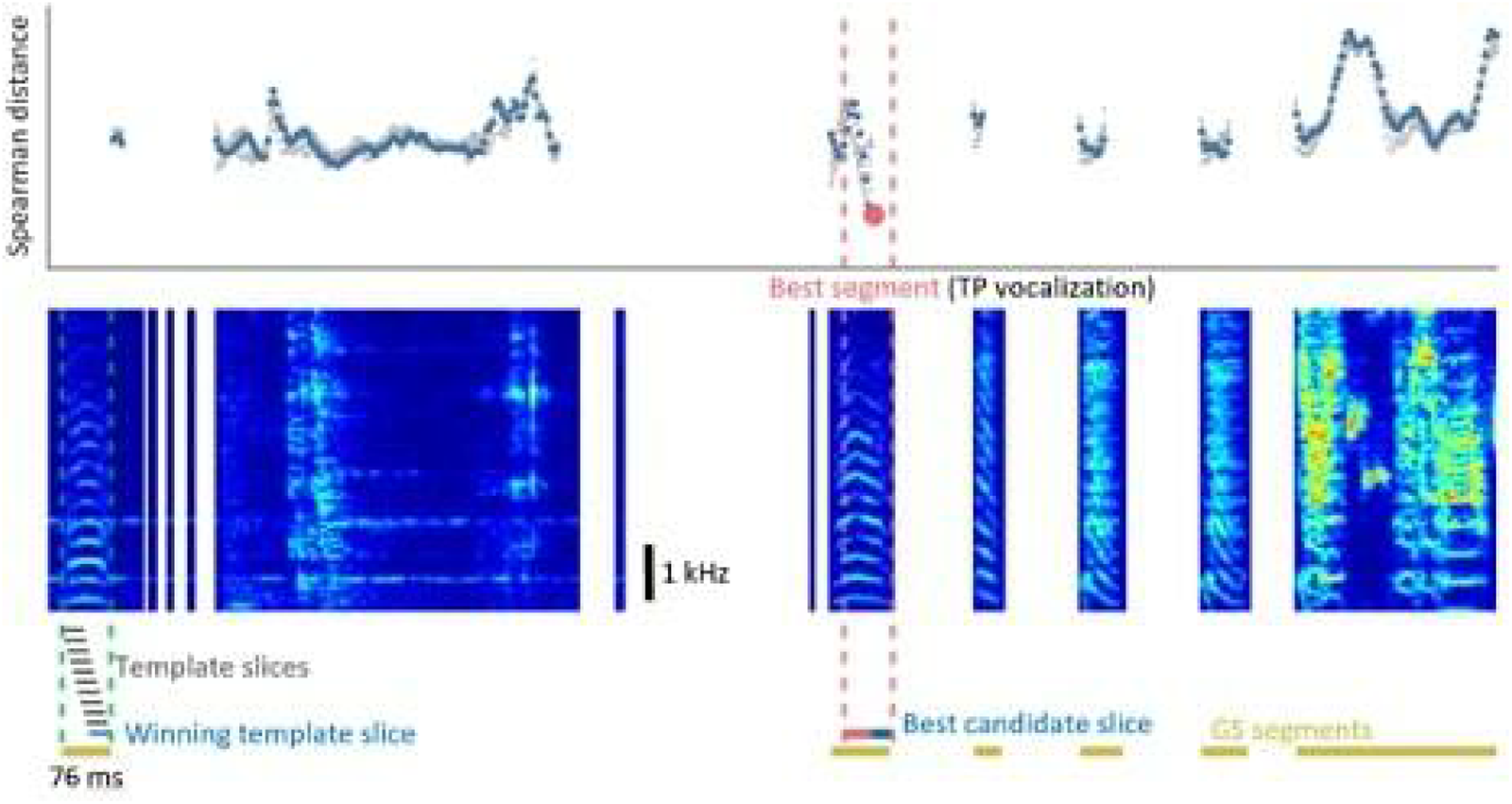
Template-based NN retrieval of vocal segments (PART approach). Shown is an example template (delimited by green dashed lines, left) that we chopped into overlapping slices (gray bars, below) of width w. For each of these slices, we computed the Spearman distances (dots, top) to candidate slices. The winning template slice (thin blue bar, bottom) and the best candidate slice (red dot, top; thick blue bar, bottom) are the ones with minimal distance to each other. From this best candidate slice, we retrieved the best segment (delimited by dashed red lines) as the sound interval that protrudes in the same way as the template relative to its winning slice. Here, this candidate is a true positive, because its relative onset (+5 columns) and offset (+1 column) are both within the accepted tolerance (*≤*5 columns) of a GS segment.

### F. Spectrogram-based distance measures

As metrics for distances *D_ij_*, we tested the Euclidean, cosine, Jaccard, and Spearman metrics using the built-in MATLAB function pdist2. Additionally, for the WHOLE approach, we evaluated earth mover’s distance (EMD) that measures the transport of sound-intensity along a single spectrogram axis: either summing EMD distances row-wise (EMDr, transport along the temporal axis) or summing column-wise (EMDc, transport along spectral axis).

### G. Performance evaluation

We evaluated the retrieval performance of our NN approaches using scores based on time bins and on sound segments:

- The time-bin based (or column-wise) score corresponds to the F1 score (the harmonic mean of precision and recall) of the inferred labels of all spectrogram column relative to the GS labels. Fig. 1 shows examples of true-positive and false-negative labels.
- The segment-wise or vocalization score (VocScore) is the F1 score of detected vocal segments. A segment is considered a true-positive (TP) vocalization if both its predicted onset and offset are within a temporal tolerance *ɛ* of the gold-standard values. This tolerance reflects the fact that even experts disagree on precise segment boundaries. Here, we have chosen a generous tolerance of *ɛ* = 5 spectrogram columns, corresponding to a generous tolerance of 20 ms on Subset 1.

## III. RESULTS

### A. A gold-standard (GS) dataset of juvenile and adult vocal segments

From a small set of template vocalizations, we performed NN retrieval of vocal segments (see Section II). We manually corrected the obtained segments to assemble a GS dataset of 53’326 vocalizations extracted from a total of 370 mins of data from zebra finches recorded at different developmental stages (Table I). We share our guidelines for manual correction that specify two decision boundaries we used to correct the segments: the decision whether there is a short silent period (gap) between two vocalizations (Fig. 5), and the distinction between vocal and non-vocal sounds (Fig. 6-7). In short, we advocate the definition of vocal segments as tight intervals of contiguous vocal activity (no gaps) (see Appendix).

**TABLE I:** Dataset of zebra finch vocal segments across 4 developmental stages. The birds’ ages are specified in days-post-hatch (dph). The last four columns specify the duration of the annotated recording (including silence and noise), the number of annotated vocalizations, the fraction of time with vocal activity (“label imbalance”, vocal/total columns; perfect balance corresponds to 0.5), and the duration range of vocalizations, respectively. The Group column refers to the recording date, i.e., the number of days (20, 10, or 0) before birds learned their baseline (BL) song (Fig. 3c).

| Developmental stage | Bird name | Sex | Hatch date | Age (dph) | Group | Annotated (mins) | Number of vocalizations | Label imbalance | Vocalization duration range (ms) |
| --- | --- | --- | --- | --- | --- | --- | --- | --- | --- |
| Adult<br>(subset 1) | g17y2 | male | 14.4.2015 | 197 |  | 84.34 | 10050 | 0.4714 | 20-656 |
|  | g4p5 | male | 28.12.2012 | 115 |  | 104.18 | 26045 | 0.5155 | 16-300 |
|  | g19o3 | male | 13.11.2015 | 154 |  | 7.72 | 2045 | 0.4238 | 20-240 |
|  | g19o10 | male | 08.11.2015 | 198 |  | 7.68 | 1998 | 0.548 | 28-400 |
| Juvenile<br>(subset 2) | R3406 | male | 29.11.2011 | 35 | -20BL | 1.27 | 139 | 0.22 | 20-357 |
|  |  |  |  | 45 | -10BL | 8.28 | 243 | 0.0486 | 9-377 |
|  |  |  |  | 55 | BL | 39.42 | 2281 | 0.1077 | 12-372 |
|  | R3428 | male | 16.12.2011 | 39 | -20BL | 7.30 | 1316 | 0.2931 | 15-514 |
|  |  |  |  | 49 | -10BL | 6.86 | 780 | 0.2496 | 12-418 |

TABLE I – *Continued from previous page*
| Developmental stage | Bird name | Sex | Hatch date | Age (dph) | Group | Annotated (mins) | Number of vocalizations | Label imbalance | Vocalization duration range (ms) |
| --- | --- | --- | --- | --- | --- | --- | --- | --- | --- |
| Juvenile<br>(subset 2) | R3428 | male | 16.12.2011 | 59 | BL | 52.19 | 4026 | 0.1862 | 23-435 |
|  | R3549 | male | 17.02.2012 | 43 | -20BL | 7.33 | 781 | 0.2411 | 15-581 |
|  |  |  |  | 53 | -10BL | 9.02 | 929 | 0.2209 | 15-438 |
|  |  |  |  | 63 | BL | 10.52 | 1068 | 0.2372 | 12-343 |
|  | R3625 | male | 13.04.2012 | 45 | -20BL | 11.67 | 728 | 0.1216 | 26-372 |
|  |  |  |  | 55 | -10BL | 7.23 | 534 | 0.1363 | 12-418 |
|  |  |  |  | 65 | BL | 4.71 | 362 | 0.1575 | 15-293 |
| All |  |  |  |  |  | 370 | 53326 |  | 9-656 |

To assess the annotation consistency, we asked a second expert to perform the same manual correction of NN-retrieved segments on a subset of data (two adults and two juveniles). We quantified expert disagreement by assessing the performance of Expert 2 relative to the GS data (Expert 1) as a reference: While the F1 score was generally high across both subsets (0.981 *±* 0.014), the VocScore fluctuated more substantially (0.923 *±* 0.046). A closer inspection revealed that the adult bird g19o3 produced pairs of rapidly following vocalizations that Expert 2 interpreted as a single vocalization, resulting in a low VocScore (F1-Score: 0.975, VocScore: 0.883), while bird g19o10 displayed no such confounding vocalization pair (F1 score: 0.992, VocScore: 0.998).

### B. Performance of nearest neighbor retrieval

We tested the two template-based vocal retrieval approaches (WHOLE and PART) on our GS dataset. The NN distance of retrieved vocalizations increased monotonically with increasing number of retrieved segments, as per definition (Fig. 3a, shown for three replicates of 50 randomly selected templates). Less trivially, the precision of retrieved vocalizations decreased with the number of retrieved vocalizations (Fig. 3a-9-10). We varied the used distance metric and the normalization strategy. We found that the Spearman distance metric performed best, particularly in juveniles, while the Euclidean metric performed worst. In juveniles also, the Jaccard metric performed better than the Cosine metric. In both adults and juveniles, both EMDs performed poorly (Fig. 3b-e). In the following, we report the performance of the Spearman metric in more detail. Using WHOLE, the Spearman distance achieved an average F1 score of 0.93 *±* 0.07 (range 0.86 to 0.98) for adults (Fig. 3b and Fig. 3d, no normalization) and an F1 score of 0.63 *±* 0.18 (range 0.23 to 0.86) for juveniles (Fig. 3b and Fig. 3e, no normalization). Using PART, the performance increased for juveniles (F1 score of 0.72 *±* 0.10, range 0.51 to 0.82) but decreased for adults (0.92 *±* 0.04, range 0.88 to 0.96), see Fig. 3c for each bird individually. This significant performance gap between adults and juveniles that we observed for the Spearman metric was also true for other metrics. The Cosine distance performed well on adults (F1-score range 0.97 to 0.81), while on juveniles it yielded low scores. Distances such as the Euclidean distance and the two

Earth Mover distances performed significantly worse than the correlation-based distances even in adults, while their respective F1 scores were close to zero in juveniles. In general, distance metrices performed significantly better in adults than in juveniles. We normalized distances in the WHOLE approach with four different strategies based on either duration or sound amplitude (see Section II). For adults, not normalizing was among the best strategies for the Spearman distance (though neither in adults nor juveniles, normalization had a large impact) and it was the worst for Earth mover’s, Jaccard, and Euclidean distances (Fig. 3d). As expected, these latter distances benefit from division by the template duration to counteract the unequal dimensions of the competing candidates. The template-wise min-max normalization worked well across distance metrics and GS data subsets (Fig. 3d,e). Taken together, NN search performed best using the PART approach on juveniles and the unnormalized WHOLE approach on adults. Across development, zebra finches can change their songs to join or to separate adjacent vocalizations (Fig. 6). To quantify errors resulting from falsely joining or separating adjacent vocalizations, we used the VocScore. The VocScore is very sensitive to segmentation errors occurring in between two vocalizations, e.g., when a syllable gap is missed, the VocScore reports a long false-positive (FP) and two short false negative (FN) vocalizations. Across both adults and juveniles, the VocScore correlated with the F1 score (Fig. 3f) and the VocScore performance was quite variable across datasets, which was due to some birds persistently producing hard-to-segment vocalization pairs. The simpler F1 score of misclassified spectrogram columns was sensitive to the number n of templates used, but surprisingly the F1 score barely improved from using more than 50 templates (Fig. 3g). The F1 score also improved with increasing slice width w (Fig. 3g), especially from the minimal width w=1 to *w* = 8. However, in juveniles, there was no additional improvement from increasing the slice width to *w* = 16 (Fig. 3g).

**FIG. 3.**
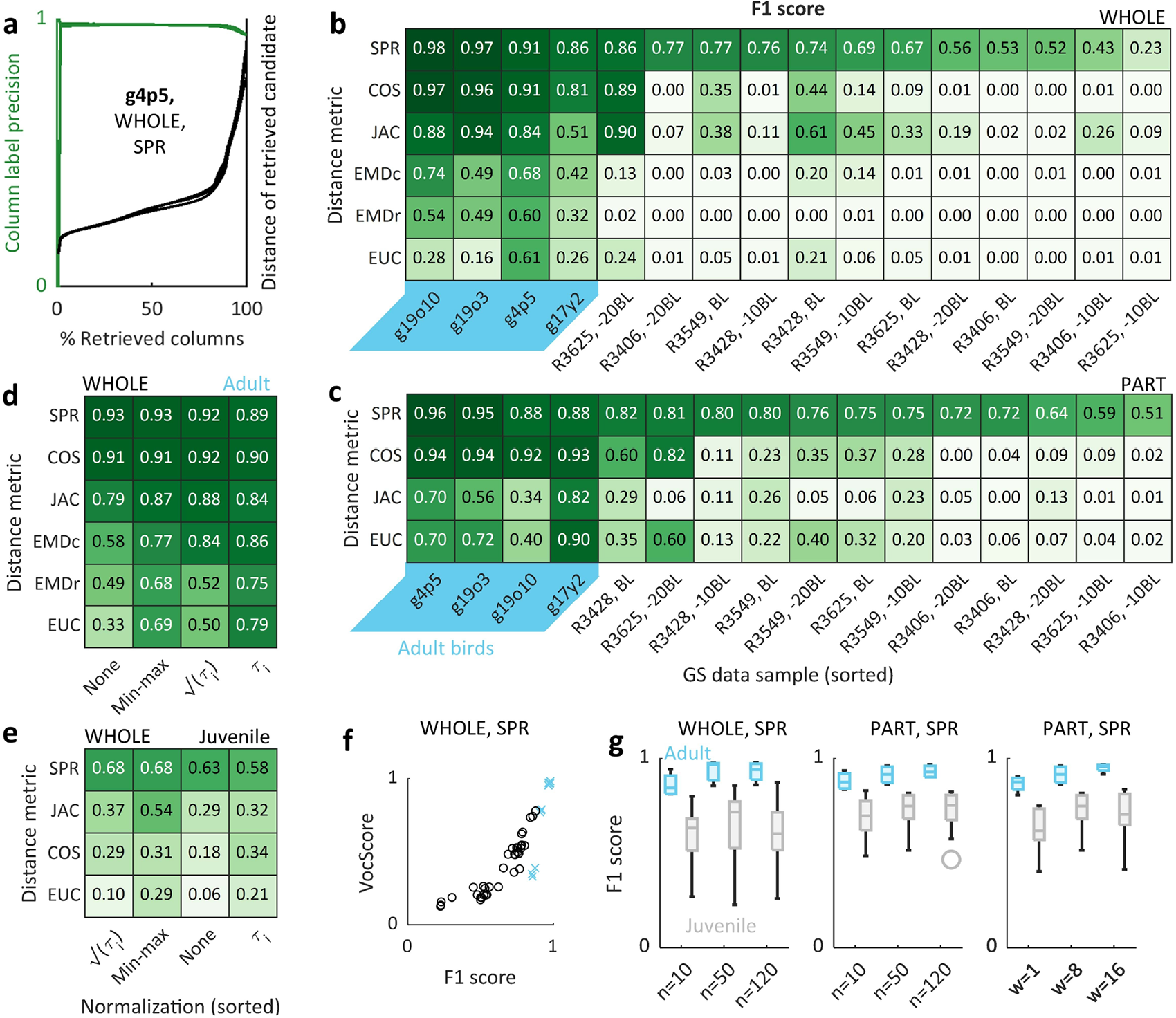
Performance of vocal segment retrieval for various distance metrics and normalization strategies. (a) The column-wise precision (green) of vocal segments gradually declined (after initial fluctuation) with increasing number of retrieved segments. We retrieved a total of N-n segments (*n* = 50 templates, *N* = 26045 GS segments, bird g4p5), corresponding to theoretical optimum of 100% of retrieved columns (x-axis). Three overlapping curves are shown for 3 replicates of 50 randomly selected templates. (b,c) Mean F1 scores (from 3 replicates of 50 random templates) across the dataset for different distance metrics, using the unnormalized WHOLE (b) or PART (c) approach (slice w=8 columns). The tables are sorted along the rows and columns to display the best performance on the top left. Abbreviations: SPR=”Spearman”, JAC=”Jaccard”, COS=”Cosine”, EMDc=”column-wise Earth mover’s distance”, EMDr=”row-wise Earth mover’s distance”, EUC=”Euclidean”. (d,e) Sorted tables of mean F1 scores (from b) of adults (d) and of juveniles (e) for the WHOLE approach, shown for different normalization strategies. (f) The relationship between F1 score and VocScore in adults (blue crosses) and juveniles (black circles), computed for the Spearman distance and using the WHOLE approach (3 replicates per sample). (g) Sensitivity analysis for the number of templates n and the slice width w, using the Spearman distance.

To investigate whether the retrieval process is hampered by some detrimental templates that excessively often retrieve false positives, we examined one retrieval replicate each in three exemplary birds, an adult and two juveniles (Fig. 4). In both birds, we found that the retrieval fractions were very non-uniform across the 50 templates (Fig. 4a-c, Figure S6, S7). In the juveniles, there were a few templates that yielded excessively low retrieval precision (large fraction of FPs). These detrimental templates had either background noises (e.g., Fig. 4b, templates “1” and “2”) or very faint harmonic extensions (e.g., Fig. 4b, template “3”). To illustrate their shortcoming, we plotted the segments retrieved by the three templates with the lowest retrieval precision in each bird (Fig. 4a-b, bottom row of spectrograms). Removing the worst three templates (searching with 47 templates only) did not increase performance in the adult (Fig. 4c), but slightly increased the performance in the juvenile (Fig. 4d). This indicates that NN search can only marginally be improved by selecting representative and clean (noise-free) templates.

**FIG. 4.**
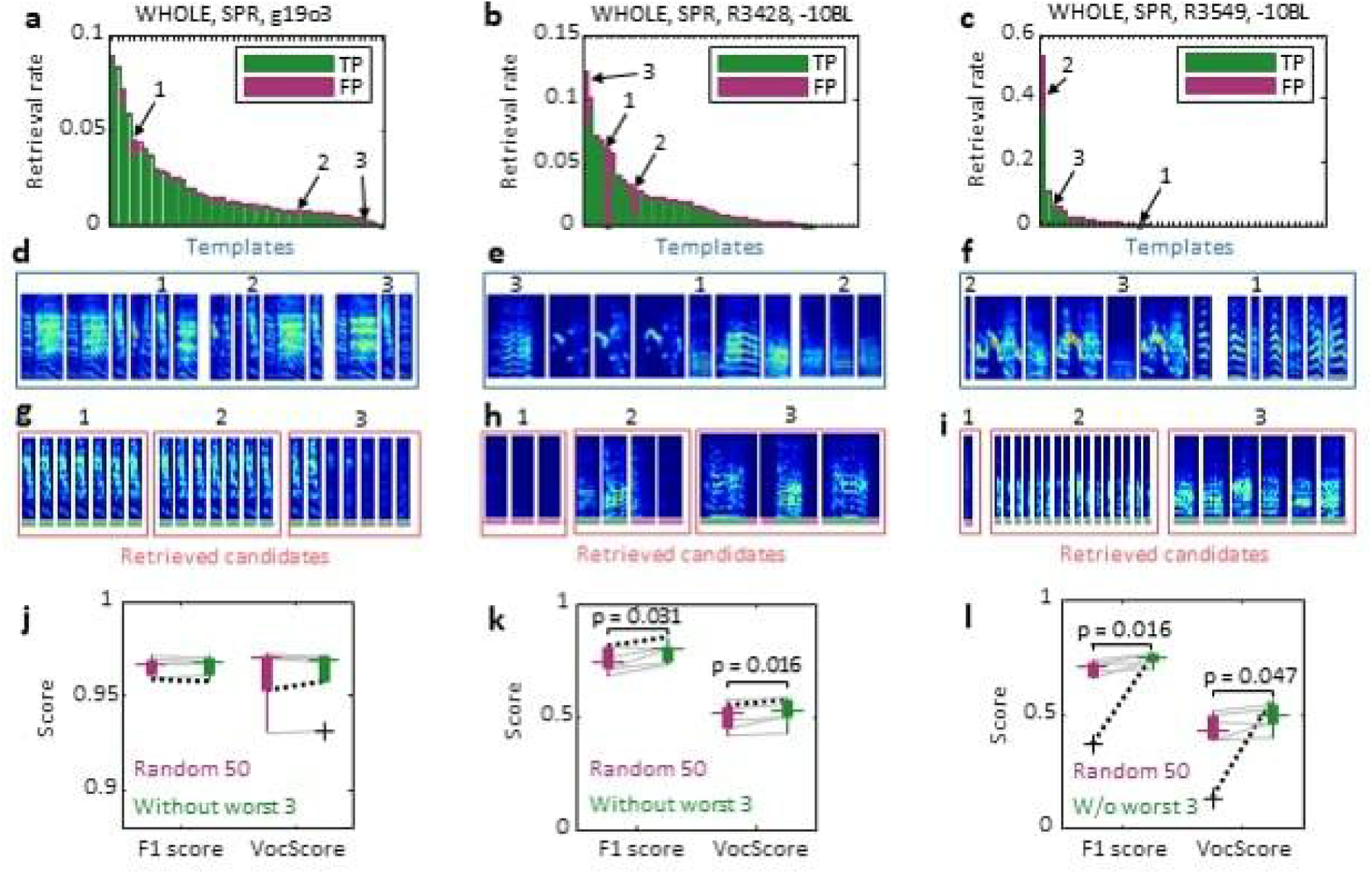
Retrieval performance is non-uniform across templates. (a,b) For an example adult (a) and two juveniles (b,c), we sorted the 50 templates (from one replicate) by the fraction of segments they retrieved (summed TP and FP retrievals). (d,e,f) For each bird, example templates are shown including the worst three (numbered 1-3). (g,h,i) Example segments retrieved by the worst three templates in each bird. (j,k,l) Performance scores (6 replicates per bird) for the initial set of random 50 templates (purple box) and for the reduced set (green box) constructed by removing the worst 3 templates. A small but significant increase in both F1 score and VocScore is observed for the juveniles (*p <* 0.05, one-sided paired-sample Wilcoxon signed rank test). The performance changes for the replicates in (a-i) are highlighted by black dotted lines (grey lines indicate changes for the remaining 5 replicates).

## IV. DISCUSSION

We have presented a simple and viable method for creating and proofreading of GS datasets of animal vocalizations. Nearest neighbor retrieval is straightforward in its application and is suitable both for extending manual annotations based on a few examples and for proofreading existing datasets. We have used NN retrieval in a 2-step process of 1) detecting vocalizations in raw sound recordings based on few labelled examples, and 2) systematic screening the remaining data for false negative samples. We evaluated NN retrieval on vocalizations from individual birds including the notoriously challenging subsongs produced during an early developmental phase. We benchmarked two NN variants and found that adult vocalizations were better retrieved using whole templates (WHOLE approach, Fig. 1) whereas juvenile vocalizations were better retrieved using template slices (PART approach, Fig. 2). We found that as few as 50 templates were sufficient for reaching plateau performance, which imposes a minimal requirement on the human effort for adopting this method. In theory, NN retrieval can be performed with as little as one single positive example. In practice, we recommend selecting clean templates and disregarding templates that contain background noises or outlier features (Fig. 4), because otherwise the noise itself becomes a target of NN retrieval. A good strategy might be to perform a two-stage search: first with stereotyped templates, then with apparent outliers. The Spearman distance outperformed the other tested metrics (Fig. 3) – especially on juvenile data. Surprisingly, the Euclidean metric, often the first choice when comparing songbird vocalizations^3,29,33,34^, exhibited the overall worst performance. That the Spearman distance outperformed the Euclidean distance on both juveniles and adults suggests that commonly used analysis methods based on the Euclidean distance^3,33^ could be improved simply by the use of Spearman distance. The finding that correlation-based metrics (including Spearman and cosine distances) outperform the Euclidean and EMD distances emphasizes the importance of discounting for vocal variability: Under the Euclidean and EMD metrics, a loud candidate vocalization will have a large distance to its softer template. Variability of sound intensity can arise from varying distances and directions of a bird to the microphone and so they should not affect retrieval. In contrast, correlation-based metrics are invariant to global changes in signal intensity (or loudness). Furthermore, correlation-based metrics work well with templates of different durations since the correlation between two vectors does not scale with the vector dimension. These results are in line with a general trend away from the Euclidean distance towards correlation-based metrics: The advantage of Spearman distance over the cosine distance is that the former captures non-linear monotonic relations^35,36^. This property is generally believed to contribute to the good performance of the Spearman distance in applications as diverse as spam email detection^37^ and indoor localization based on received Wi-Fi signal strength^38^. We see the strength of NN retrieval in proofreading the predictions generated by other systems, in particular when labelled data are scarce. By contrast, when labelled data are abundant, NN retrieval is unlikely going to be competitive with state-of-art approaches for birdsong segmentation such as deep neural networks^7,8^. The main disadvantage of NN retrieval (e.g. compared to neural networks), is that the computational cost scales with the number of labelled examples, although workarounds could be to sub-sample or summarize the templates using for example k-means clustering. Very large datasets are amenable to NN retrieval by virtue of powerful methods for approximative NN retrieval^22–25^. Therefore, there is no fundamental barrier for scaling up this method. We benchmarked NN retrieval on vocal segmentation, which is a task that is feasible in both adults and juveniles and allows for comparison of performance across age. In adults with their stereotyped repertoire, it is possible to target retrieval to renditions of specific syllable types rather than any vocalization from the repertoire. Coincidentally, we used such type-specific retrieval to generate the GS annotations for adults. In practice, we found that best performance is achieved when first searching for renditions of long vocalization types and then successively for shorter types. Such a hierarchical retrieval strategy avoids confounds from repeated notes among syllables in adult zebra finch song^39^, which may also be the reason for the lower performance of PART in adults compared to WHOLE. By contrast, the reason why for juveniles, PART seems to work better than WHOLE could be that on a larger time scale juveniles have no repeating vocal units — thus, if we model their vocalizations as random vectors then these are all far from each other since in large spaces, random pairs of vectors tend to be orthogonal to each other. Our retrieval approach (in particular the WHOLE approach) suffers from inflexibility of segment durations, namely that the retrieved segments must exhibit the same durations as the templates. Therefore, WHOLE will struggle to find the overall shortest vocalization performed by an animal. One possible approach to overcome this limitation is to use dynamic time warping^29^ as a means to create artificially short templates, thereby increasing the number and diversity of templates. NN retrieval is attractive because it controls for out-of-distribution detection with a well-defined and interpretable distance measure. NN retrieval shifts the challenge of modeling vocalizations to the challenges of identifying a good metric. We tested only a set of well-known metrics here, but in follow-up work it may be worthwhile train custom metrics on the same retrieval task to learn to optimally account for natural variability. Metrics can be learned from embeddings and the approach of computing embeddings in a self-supervised manner^40^ is getting more popular also in sound processing^41^, in particular speech^42,43^. The role of NN search we foresee in future work is to assist in creation of vocal annotations and in proofreading automated annotations produced by trained systems. One promising idea is to develop human-in-the loop iterative procedures of labelling, training, searching, and fine-tuning of machine-learning systems. Our expert-curated dataset of annotated individual vocal repertoires counts more than 50’000 vocalizations from 8 zebra finches. We release this dataset so that data-hungry deep learning systems for large scale vocal analysis can be trained and evaluated. To make our work reproducible, we also share our segmentation guidelines as illustrations of the manual annotation challenges and of our chosen decision boundaries (see Appendix). We hope that our annotation guidelines will help to standardize vocal annotation tasks and so promote comparative work across species.

## DATA AVAILABILITY

We will release our dataset (Table I) upon publication of our work in a peer-reviewed journal.

## AUTHOR CONTRIBUTION

RHRH, TT, SR, and XH contributed to the conceptualization of the study. ATZ conducted experiments of Subset 1. TT, RHRH, XH, and AM contributed to data annotation. TT and ATZ curated the dataset for release. TT and RHRH implemented the retrieval algorithms. TT, RHRH, XH, SR, and KL were involved in data analysis. TT, ATZ and RHRH wrote the manuscript. SR, AM, KL and MB provided feedback on the manuscript.

## ACKNOWLEDGEMENT

We thank Dina Lipkind for making her previously published data (Subset 1) available for post-hoc analyses. Experimental procedures involved in Subset 1 of the GS dataset were approved by the Veterinary Office of the Canton of Zurich.

## APPENDIX

## 1 Vocal segmentation conventions for microphone recordings of single birds

Vocal signals tend to arise from discrete acoustic units, which is a characteristic shared across the polymorphic landscape of vocalizing species^44,45^. Animal studies in monkeys, dogs, chicken, and songbirds have shown that animal calls can be used to communicate semantic meaningful information such as detection of predators, discovery of food, or attraction of mates^46–55^. Nevertheless, the functions of animal vocalizations are generally unknown for most calls and species^44,56^. To advance our understanding of vocal communication in animals, we need to study large and well-annotated data sets. Here we address the problem of how to segment audio recordings of a given species. The segmentation problem is to distinguish the times at which an animal vocalizes from the times at which it does not. One of the simplest methods of segmenting vocalizations from continuous recordings is to consider sound amplitude and to define as vocalizations all sounds that are above a given threshold. However, this procedure will misclassify certain noises as vocalizations, which is why more refined approaches are needed that potentially make use of the statistics of the individual^33^. In the extreme case, we need to inspect every single potential vocalization and decide based on expert knowledge where to cut the dividing line between vocalization and noise.

To standardize the segmentation task, we have created this set of guidelines based on two decisions boundaries for a vocalization:

- The decision whether there is a silent period between two sounds, which we take by inspecting spectrograms (Fig. 5, left).
- The decision whether a sound is vocal or non-vocal (Fig. 5, right; Fig. 6-7).

Birds, especially when young, tend to vary the gaps between vocalizations. An example is shown in Fig. 5 (yellow dotted box): This sequence of three vocal elements looks like a precursor of syllable C that the juvenile tries to imitate, but they appear with sufficiently large gaps, which is why we sometimes classify them as 3 distinct syllables. Thus for (a) we infer a gap where we can visually detect one, irrespective of other singing attempts in the animal. The second decision boundary (b) is harder to define universally from single-microphone recordings, ideally we would like to have simultaneous recordings from the trachea to measure sounds and air flow there. In practice, it is a human expert, who judges whether a sound is vocal or non-vocal by listening to examples and inspecting the corresponding spectrograms. Again, this task is relatively simple for highly stereotyped vocalizations, but more difficult for faint, short and variable vocalizations in juveniles (Fig. 5, right; Fig. 6, left, Fig. 7). A special case consists of faint sounds (usually at around 6kHz) that frequently occur after (or, less frequently, before) vocalizations (Fig. 2, left). We consider them to be inhalation sounds^33,57^ and exclude them from the vocal dataset (default setting).

**FIG. 5.**
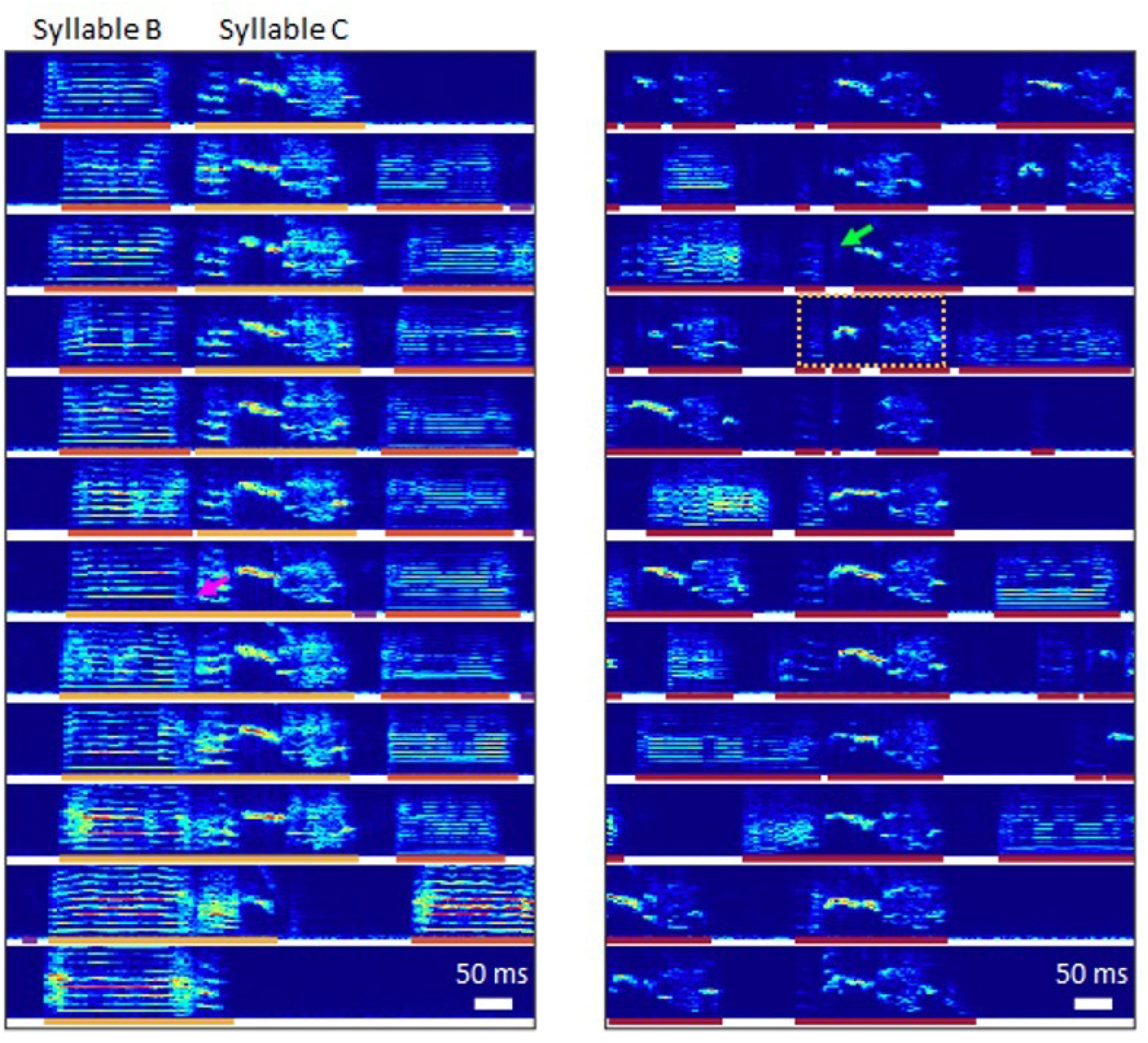
Definition of vocal segments as continuous intervals of vocal activity. (left) Zebra finch song examples at 59 day-post-hatch, aligned to notes that resemble the beginning of syllable C. At this stage, syllable C is surrounded by clear gaps most of the time (top 6 examples). However, in a minority of cases, no silent gap is visible between the preceding syllable B and the first note of syllable C (bottom 6 examples, boundary case indicated with magenta arrow). Gold-standard segmentation labels of syllable-C-notes (yellow) and of other vocalizations (orange, purple) are indicated by bars below the spectrograms. (right) Vocalizations recorded at 49 day-post-hatch (red bars), aligned to examples that resemble syllable C. Short noisy sounds within syllable precursors (green arrow) have not been classified as vocal activity based on isolated visual inspection, but likely would be, if the context would be taken into account. The yellow dotted box marks three vocal elements that could potentially be interpreted as a unitary precursor of syllable C, if the developmental endpoint were to be taken into account. Bars as on the left.

**FIG. 6.**
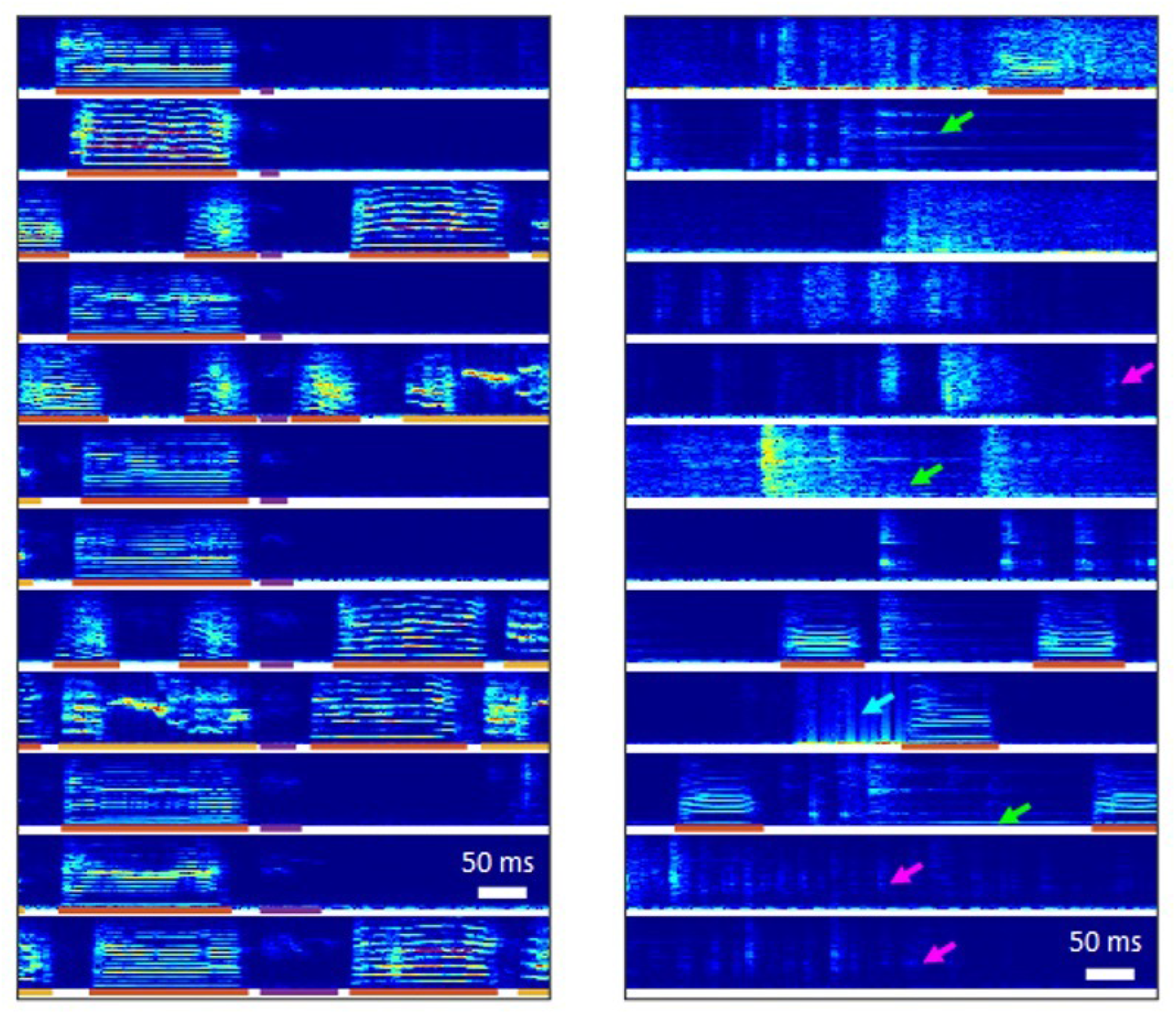
Decision-boundary between vocal and non-vocal sounds. (left) Spectrogram examples of putative inhalation sounds (indicated with purple bars) observed in a zebra finch at 59 day-post-hatch (excluded in the gold standard by default). (right) Examples of non-vocal noises which may include prominent tones (green arrows), wide-band noise (blue arrows), or very faint signals (magenta arrows).

**FIG. 7.**
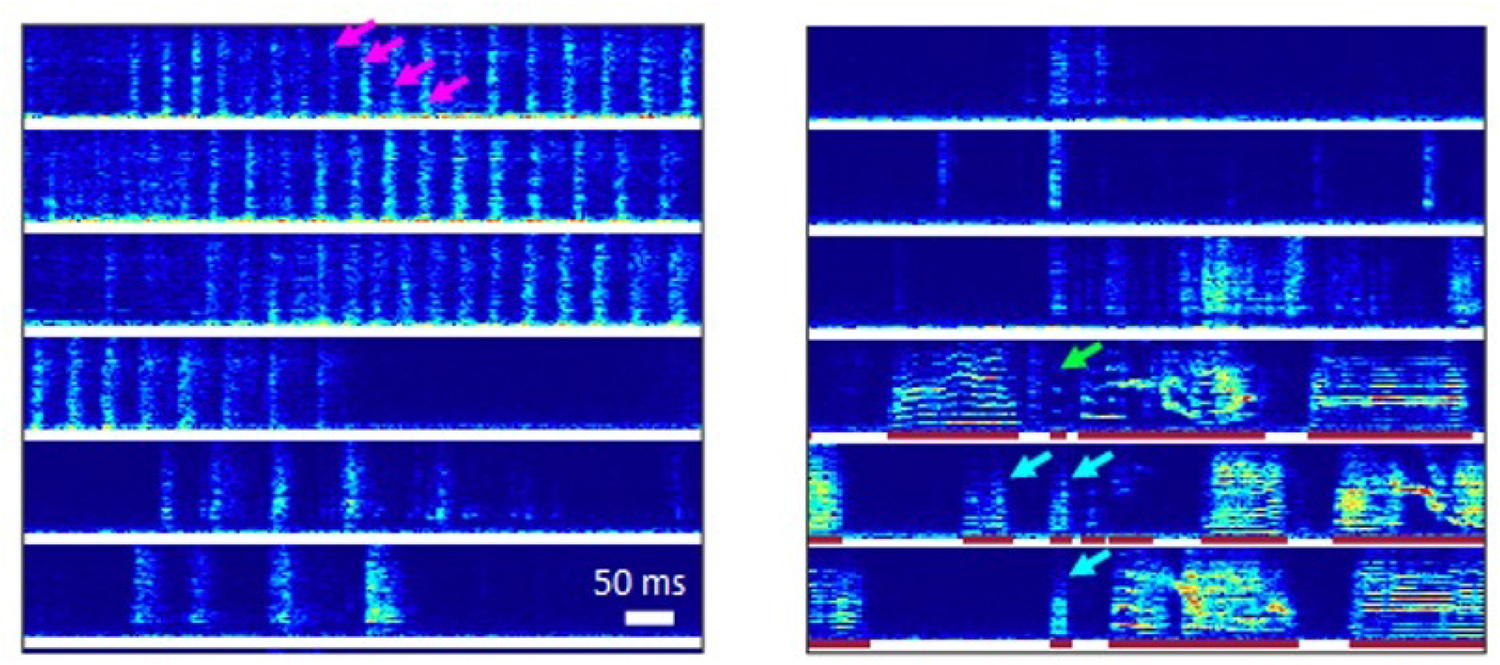
Detailed decision-boundary between vocal sounds and wing flaps. Spectrogram examples short noises. Wing flaps are easy to detect on spectrograms when occurring in serial repetition (i.e., when the bird is flying; magenta arrows). For short sounds, indicators of vocal activity can be harmonics (green arrow) or a strong skew in the spectral density towards certain frequencies (low frequency sounds indicated with blue arrows).

## 2. Analysis of an open dataset

A recent publication^12^ includes a large dataset of vocal segments from 5 zebra finches. According to the data documentation, the segmentation was performed using a sound-amplitude based method that included some hand tuning. Although we found the published segmentation results to be valuable, they were insufficient to qualify as gold standard, due to the existence of false negatives and inaccurate segment boundaries Fig. 8.

**FIG. 8.**
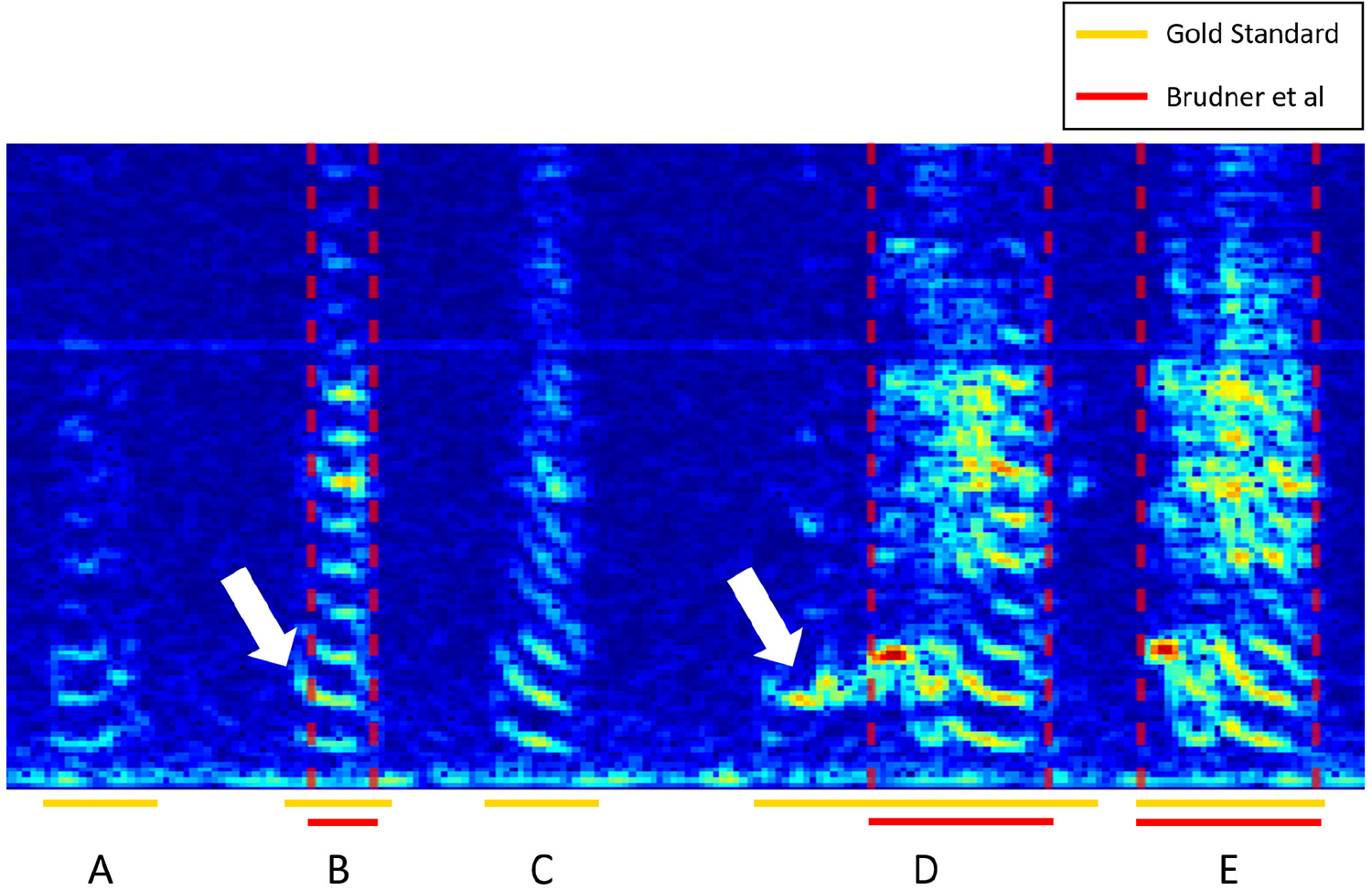
Example segmentation inaccuracies of the^12^ dataset. The published segments (red horizontal bars) deviate from the (gold-standard) manual annotations (gold horizontal bars) in terms of a false negative sample (Syllables A and C) and in terms of inaccurate segment boundaries (white arrows).

## 3. Discussion

The examples we provided illustrate our decision boundaries and the difficulties with segmentation approaches. In summary, we advocate the definition of vocal segments as tightly restricted intervals of continuous vocal activity. These segments should be defined independently from functional considerations. How to extract functional units from vocal segments is an open question, the answer may depend on whether the vocal units are assessed in the domain of perception (receiver) or production (sender). Still, it is regarded as ideal to validate chosen segmentations based on the functional roles of the vocal signals^44,56,58^. However, recent work in songbirds suggests that “syllables may not be perceptual units for songbirds as opposed to common assumption”^59^.

**FIG. 9.**
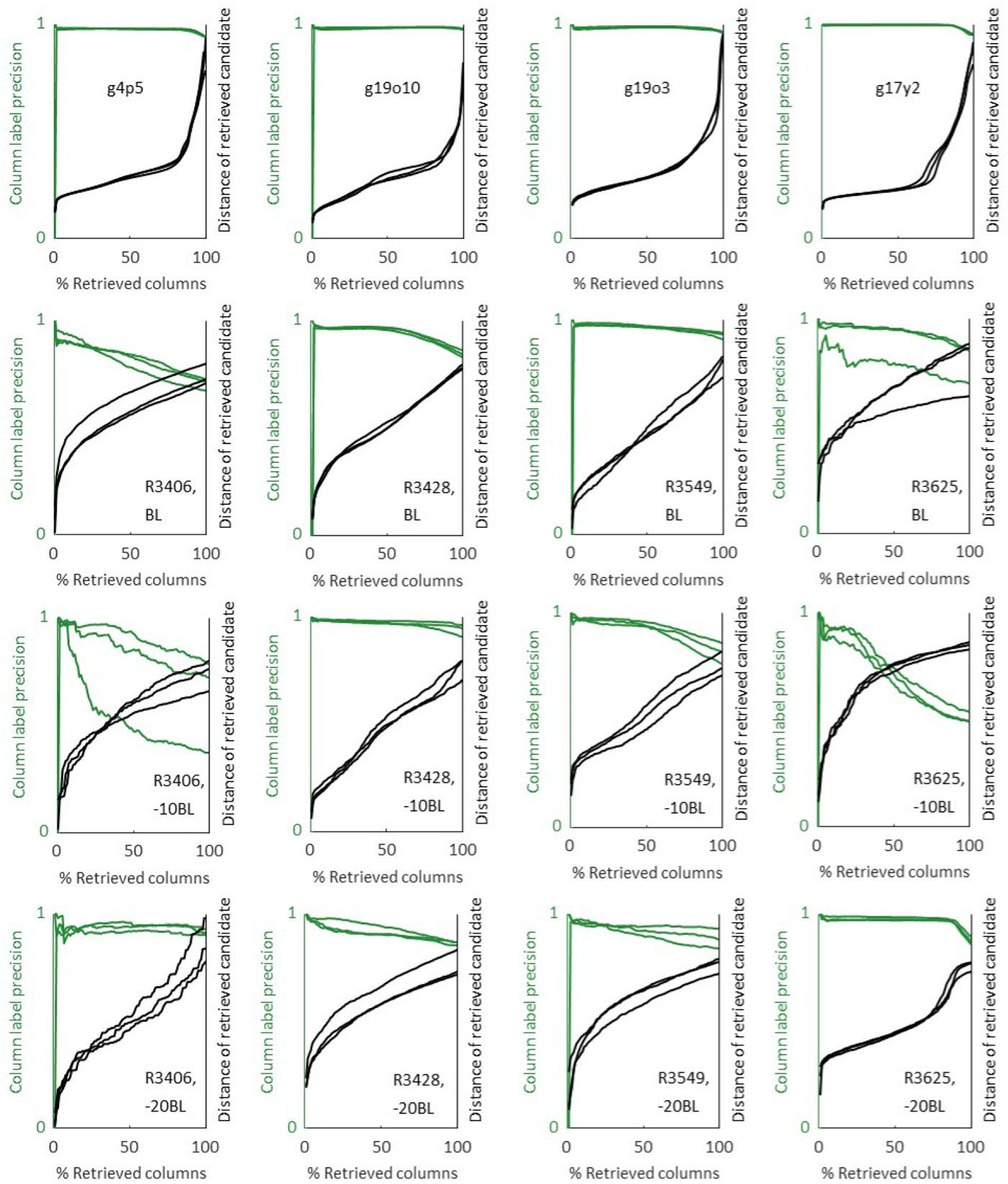
Extended set of precision and distance curves as a function of retrieval progression, using the WHOLE approach (replicated for all birds). The top row shows adult birds, while the subsequent rows show juveniles at different ages relative to baseline. See Figure 3a for a detailed description.

**FIG. 10.**
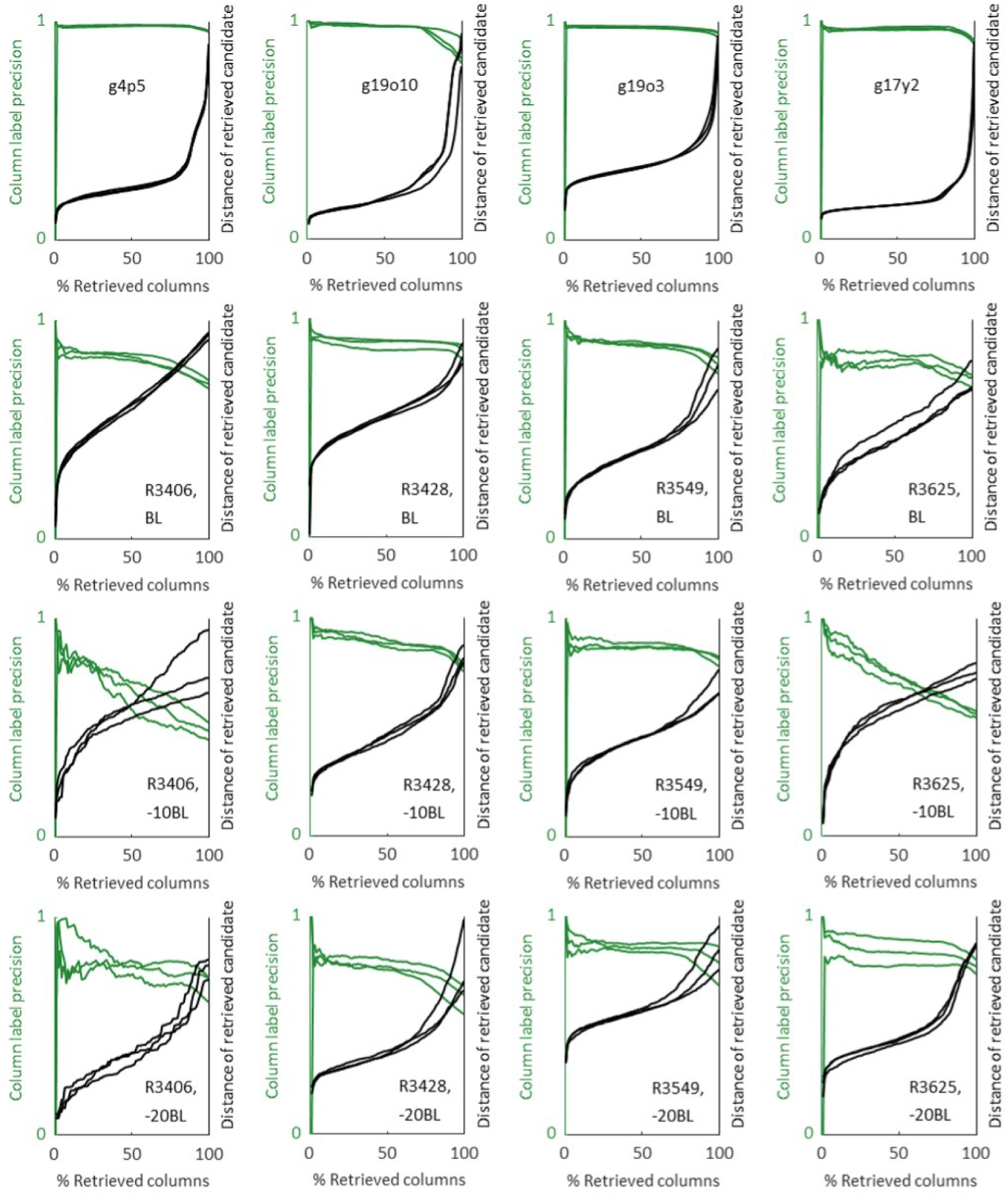
Extended set of precision and distance curves as a function of retrieval progression, using the PART approach (replicated for all birds). The top row shows adult birds, while the subsequent rows show juveniles at different ages relative to baseline. See Figure 3a for a detailed description.

**FIG. 11.**
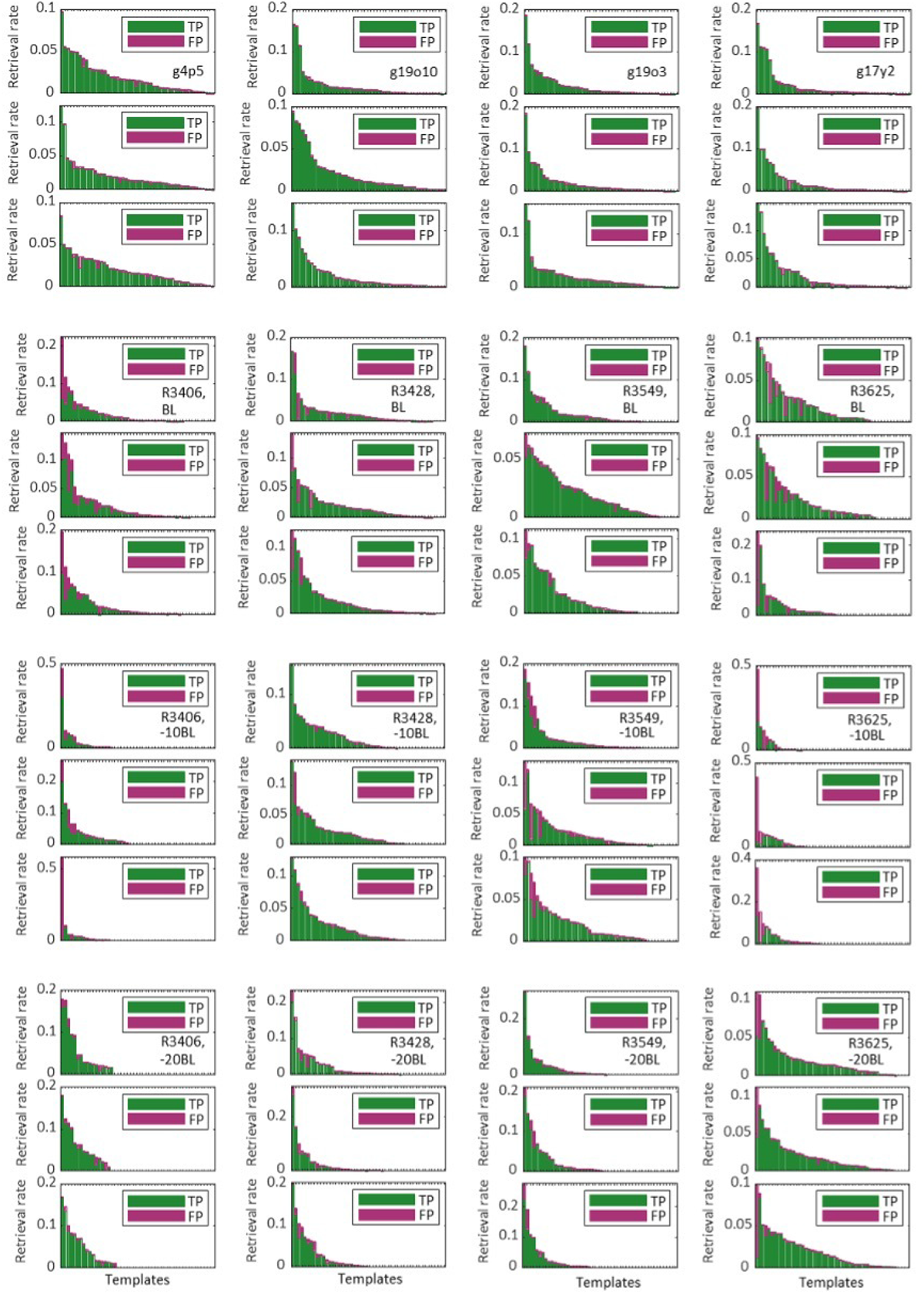
Extended set of histograms of retrieval rates across templates, using the WHOLE approach (3 retrieval replicates for each bird). The top row (consisting of 3 panels for each retrieval replicate) shows adult birds, while the subsequent rows show juveniles at different ages relative to baseline. See Fig. 4a-c for a detailed description.

**FIG. 12.**
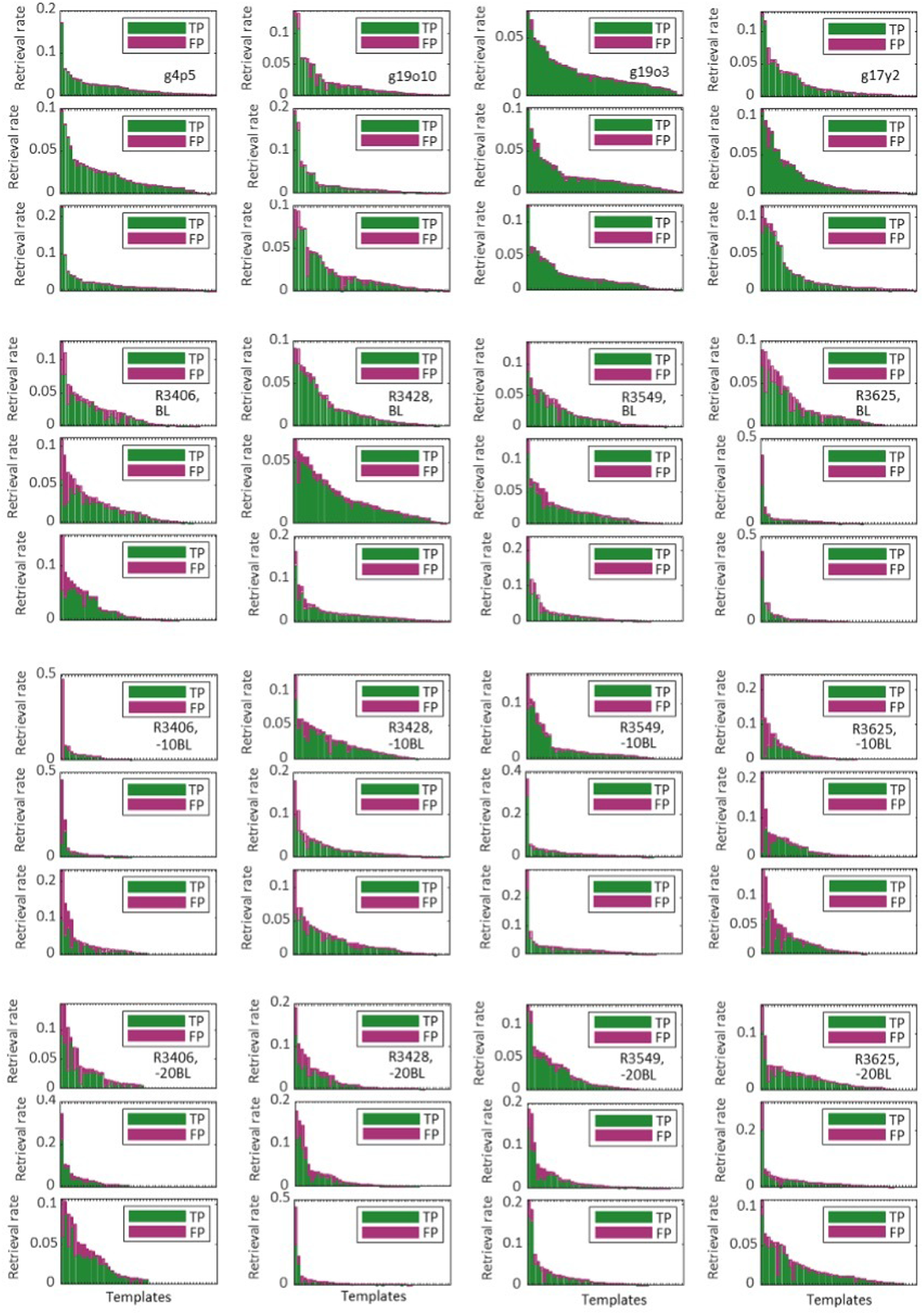
Extended set of histograms of retrieval rates across templates, using the PART approach (3 retrieval replicates for each bird). The top row (consisting of 3 panels for each retrieval replicate) shows adult birds, while the subsequent rows show juveniles at different ages relative to baseline. See Fig. 4a-c for a detailed description

